# Multiomics-based assessment of 2D and 3D human iPSC-cardiomyocyte models of insulin resistance demonstrate metabolic and contractile dysfunction that recapitulates diabetic cardiomyopathy

**DOI:** 10.1101/2024.11.20.624467

**Authors:** Ryan D. Carter, Ujang Purnama, Marcos Castro-Guarda, Claudia N. Montes-Aparicio, Anandhakumar Chandran, Richard Mbasu, Maxwell Ruby, Charlotte Daly, Francesca M. Buffa, Lisa C. Heather, Carolyn A. Carr

**Author notes:** Denotes joint first author (Ryan D. Carter, Ujang Purnama). Denotes joint last author (Lisa C. Heather, Carolyn A. Carr). Address for Correspondence: Associate Professor Lisa Heather Department of Physiology, Anatomy and Genetics University of Oxford Oxford United Kingdom.

## Abstract

In type II diabetes (T2DM), the heart is exposed to hyperglycaemia, hyperlipidaemia, and hyperinsulinaemia, leading to insulin resistance and metabolic dysfunction, culminating in diabetic cardiomyopathy (DbCM). Human-centric models of DbCM are needed to provide mechanistic insights and therapeutic targets in a translationally relevant setting. We hypothesised that culturing human-induced pluripotent stem cell-derived cardiomyocytes (hiPSC-CMs) in an “insulin resistance” (IR) media, and assessing this using a systems biology approach, would offer a comprehensive evaluation of dysregulated pathways, establishing their suitability as a model of DbCM.

Culturing hiPSC-CMs in 2D or 3D as engineered heart tissue (EHT) in IR media induced insulin resistance and activated numerous pathways implicated in DbCM, including metabolic remodelling, mitochondrial dysfunction, extracellular matrix remodelling, and endoplasmic reticulum stress. Pathways involved in fatty acid oxidation were upregulated, while those involved in glucose metabolism were downregulated, which was validated using radioisotope flux measurements. Adaptation to hypoxia, a key component of post-ischaemic remodelling, was blunted in the 2D IR hiPSC-CMs. Combining proteomic and transcriptomic analyses in the IR 3D EHT revealed significant enrichment of DbCM pathways, with subnetworks enriched for several metabolic and diabetes-related pathways. Additionally, IR 3D EHT displayed impaired relaxation, mimicking the diastolic dysfunction observed in T2DM patients.

In conclusion, culturing hiPSC-CM in 2D or 3D in an IR media activates multiple mechanisms implicated in the development of DbCM, with IR 3D EHT also recapitulating the diastolic dysfunction present in patients with T2DM.

## Introduction

Type 2 diabetes mellitus (T2DM) is characterised by diminished systemic metabolic homeostasis, affecting many organs throughout the body. T2DM is strongly linked to an increased risk of cardiovascular disease (CVD), which is the leading cause of death among T2DM patients^1^. The relationship between T2DM and CVD is complex, with multiple risk factors such as hypertension, coronary artery disease and peripheral arterial disease developing concurrently^2^. However, T2DM can also independently affect cardiomyocyte structure and function, known as diabetic cardiomyopathy (DbCM)^3^, with hallmarks of DbCM often present before evidence of symptomatic cardiovascular disease. These hallmarks of DbCM include abnormal substrate metabolism, fibrosis, diastolic dysfunction and hypertrophy^4^.

In T2DM, the heart is exposed to hyperglycaemia, hyperlipidaemia, and hyperinsulinaemia, which impact substrate uptake and utilisation. This environment drives metabolic derangements in the heart, characterised by increased dependence on fatty acid metabolism and decreased preference for glucose metabolism^5^, contributing to cardiac insulin resistance (IR) development. This shift results in lipid accumulation, mitochondrial dysfunction, endoplasmic stress and cardiac remodelling^6–8^. These metabolic derangements, alongside inflammation, dysregulated calcium handling, extracellular matrix remodelling, and endothelial dysfunction are critical mediators in the development of DbCM^4^.

Although up to 70% of T2DM patients exhibit hallmarks of DbCM^9^, available treatments are limited, and there is an unmet need for preclinical models to elucidate pathology and develop therapeutics. Animal models of T2DM and IR have been instrumental in uncovering fundamental DbCM mechanisms^10^. However, animal models have different responses to therapeutics than humans, which may not be predictive for clinical trials. Human heart tissue and cardiomyocytes can offer more human-relevant models; however, they are difficult to obtain (particularly healthy control tissue), hard to maintain in culture, and heterogeneous. A human-centric model of DbCM is needed that can be used alongside animal studies and patient studies to provide mechanistic and therapeutic insights in a human-relevant setting. Human induced pluripotent stem cell-derived cardiomyocytes (hiPSC-CMs) provide an alternative model that is more human-centric, readily available, and has fewer ethical considerations.

Although hiPSC-CMs have been used extensively in modelling monogenic diseases and regenerative medicine, using them to study complex multifactorial, lifestyle-related diseases like T2DM has been minimally explored^10^. Drawnel *et al.* demonstrated similar changes in calcium handling and beat rate irregularity when comparing hiPSC-CMs donated from T2DM donors and control hiPSC-CMs cultured in a diabetogenic media; thus, mimicking the diabetogenic environment is sufficient to recapitulate the disease phenotype^11^. Some studies have manipulated individual components of the diabetogenic environment by culturing hiPSC-CMs in either high glucose or fatty acid concentrations, with studies recapitulating specific aspects of IR^12,13^. One potential issue is that 2D cultured hiPSC-CMs are inherently immature, whereas culturing in 3D as engineered heart tissue (EHT) results in a more mature phenotype than the 2D cells, as demonstrated by increased mitochondrial number, sarcomere organisation and metabolic maturation towards a more oxidative phenotype^14,15^. This metabolic maturity could be necessary when modelling a metabolic disease like T2DM; a question that has not been addressed. However, when compared with 2D hiPSC-CMs, generating 3D EHT is a more time-consuming and expensive process.

A systems biology approach is needed to determine if this complex multifactorial disease of DbCM can be modelled in hiPSC-CMs, and if it is sufficient to do this in 2D or if it is necessary to carry this out in 3D EHT. Here, we have developed a protocol to induce IR in hiPSC-CMs using media that recapitulates the hyperglycaemia, hyperlipidaemia and hyperinsulinemia found in patients with T2DM. We have utilised omics-based analyses to evaluate phenotypic changes at a systems level, assessing the efficacy of 2D and 3D hiPSC-CM models in capturing the DbCM phenotype. We demonstrate that culturing hiPSC-CM in IR media induces many of the hallmarks of DbCM, including a metabolic shift away from glucose towards fatty acid metabolism, endoplasmic reticulum stress, extracellular matrix remodelling and abnormal relaxation. We demonstrate that there are advantages and disadvantages of modelling IR in 2D vs. 3D, dependent on the feature of DbCM of interest to the investigator.

## Methods

### hiPSC-CM differentiation

IMR90 human inducible pluripotent stem cells (hiPSCs) were differentiated into hiPSC cardiomyocytes (hiPSC-CM) via the temporal modulation of Wnt signalling protocol^16^. Before differentiation, hiPSCs were grown to confluency in TeSR-E8 media (StemCell Technologies) on matrigel-coated plates (Corning). Confluent hiPSCs were differentiated into hiPSC-CMs over 15 days, as previously described^14,17^. Cells were cultured on Reduced Growth Factor Matrigel-coated plates in RB-media (RPMI media (ThermoFisher) supplemented with 1% B27 minus insulin (ThermoFisher)) and the GSK3 inhibitor CHIR99021 (Tocris, 6 µM) for days zero and one. The following day, cells were cultured in RB-only. On day three, this media was swapped for RB-media supplemented with Wnt-C59 (Tocris, 2.5 µM). On day five, cells were again changed back to RB-only, replenished every two days until day eleven. To metabolically select CMs over adjacent lineages, the media was replenished with RB-without glucose on days eleven and thirteen. Finally, on day fifteen, the media was replaced with RB+ Media (RPMI supplemented with 1% B27 complete (ThermoFisher)).

### Generation of 3D Engineered Heart Tissue

The generation of engineered heart tissue (EHT) was adapted from previously published protocols^18^. Agarose (Invitrogen, 2%) was dissolved in PBS to create a sterile moulding solution, which was warmed and pipetted into 24-well plates containing Teflon spacers (EHT Technologies) that defined the negative space for casting. The spacers were removed when cooled, and silicon posts (EHT Technologies) were added as anchorage sites for EHT formation. Stock solutions were prepared of 10xDMEM (1.3g DMEM powder (Gibco) in 10 ml sterile water); supplemented 2xDMEM (20% horse serum (ThermoFisher), 2% Penicillin/Streptomycin (Gibco), 20% 10× DMEM in sterile water); Y-27632 rock inhibitor (Merck, 9 mM); aprotinin (Sigma Aldrich, 33 mg/ml); fibrinogen (Sigma Aldrich, 200 mg/ml in 0.9% NaCl) with aprotinin stock added to a concentration of 0.1 mg/ml); and human insulin (Merck, 10 mg/ml). Following differentiation, hiPSC-CMs were dissociated using TrypLE Express Enzyme (Gibco) at 37°C for 8 minutes. Approximately 1 x 10^6^ dissociated cells per EHT were added to a master mix containing 94.56 µL hydration media (DMEM containing 10% FBS, 1% penicillin/streptomycin, 1% L-glutamine (Life Technologies)), 6.68 µl of supplemented 2x DMEM solution, 0.12 µL rock inhibitor stock, 12 µL reduced growth factor Matrigel (Corning, 354230), 3.04 µl of the combined solution of fibrinogen and aprotinin. To this master mix, 3.6 µL of thrombin (Biopur, 100 U/mL) was added immediately before transferring to the agarose mould and incubating for 90 minutes. An additional 500 µL of hydration media was added before incubating for 30 minutes. Once set, the EHTs were transferred to 24-well plates and cultured in EHT media comprising 20% horse serum, 1% penicillin/streptomycin, 0.1% aprotinin stock and 0.1% of insulin stock in DMEM (1g/L glucose).

### 2D and 3D hiPSC-CM maturation protocol

Once beating, the 2D hiPSC-CMs and 3D EHTs were moved into maturation media for seven days. The maturation media comprises DMEM (Sigma Aldrich, 1 g/L glucose) supplemented with 10% horse serum, 5 µg/mL vitamin B12 (Merck), 0.5 mM ascorbic acid (Merck), 0.84 uM biotin (Merck), 1.7 µM insulin, 80 µM oleic acid: BSA (2:1, Sigma Aldrich), 1x MEM non-essential amino acids, and 0.1X penicillin-streptomycin.

### Inducing Insulin Resistance in hiPSC-CM and hypoxic exposure

Insulin resistance was induced using a six-day protocol by culturing under 2 different culture media, collectively referred to as the IR media^17^. Mature hiPSC-CMs or EHTs were moved to glucose-free insulin-resistance media, consisting of DMEM no glucose (Gibco), supplemented with 0.3 mM palmitic acid:BSA (bound 6:1, Sigma Aldrich); 1.7 µM insulin (Sigma Aldrich); 1X nonessential amino acids (Gibco), 0.5 mM ascorbic acid (Sigma); 5 µg/mL vitamin B12 (Sigma); 0.84 µM biotin (Sigma); 0.1X penicillin/streptomycin (Sigma); and 10% heat-inactivated horse serum for 3 days. For the following 3 days, hiPSC-CMs were switched to the glucose-enriched insulin-resistance media, like the previous but containing 12 mM glucose and 3.4 µM insulin. Control cells were cultured for six days in maturation media.

For one set of experiments, hiPSC-CM were cultured in high palmitate media consisting of palmitate bound 6:1 with BSA and diluted to 0.4mM in DMEM (1g glucose/L). The media also contained 5 µg/mL vitamin B12, 0.5 mM ascorbic acid, 0.84 µM biotin and 1x nonessential amino acids to support cellular functions. For the controls, the same media was used without palmitate addition. For hypoxia experiments, 2D hiPSC-CM were transferred to a hypoxic incubator for 16 hours (2% O_2_, 5% CO_2_ at 37°C) and compared with cells cultured in normoxia (21% O_2_, 5% CO_2_ at 37°C).

### RNA isolation and quantitative PCR

Cells were washed in PBS, lysed using the RNAeasy kit QIAzol lysis agent, scraped, and homogenised using needles before being flash-frozen in liquid nitrogen. RNA was extracted from the lysates using the Qiagen RNeasy Mini kit following the manufacturer’s protocol. For relative quantitative comparative PCR (qPCR), cDNA was synthesised from the RNA using a high-capacity RNA-to-cDNA kit (Thermo Fisher Scientific), and qPCR was carried out using a Step-One Plus Real-Time PCR system using Power SYBR Green PCR Master Mix (primer sequences in Supplementary Table). Relative gene expression was calculated with the 2^−ΔΔCT^ method ^19^ and normalised to the housekeeper gene ubiquitin C.

### Transcriptomics mapping, quantification, and differential expression analysis

mRNA from 2D hiPSC-CM and EHT samples was sequenced using Illumina protocols, with data processed via custom Snakemake pipelines^20^. For hiPSC-CM, sequencing involved a stranded paired 150-bp poly-A enrichment protocol by Novogene, while EHT sequencing used Illumina 3’ quantseq in collaboration with the Novo Nordisk Research Centre Oxford (NNRCO). Quality control was performed with FastQC, trimming with cutadapt^21^ to remove adaptors and low-quality sequences. Transcript quantification for hiPSC-CM was executed with Salmon^22^ using decoys and a gentrome indexed from the gencode v44 GRCh38, and reads were processed with a mapping rate of approximately 80%. For EHTs, STAR^23^ was used to map reads with similar efficiency, using an index from the Ensembl human genome version 109 GRCh38.

Differential expression analysis was conducted using the DESeq2 pipeline^24^. Lowly expressed genes were filtered out if they had fewer than ten counts in at least five samples. Counts were normalised for exploratory analysis and visualisation using the variance stabilisation transformation (VST), which corrects for mean-variance correlation and differences in library sizes and transforms counts to a ‘log-like’ scale^25^. Principal component analysis was performed using the plotPCA function, and differential expression was analysed using the specified differential testing designs. Log fold changes were adjusted using the apeglm method for more accurate effect size estimation^26^. Significance thresholds were set at p < 0.05 with Benjamini Hochberg correction. For EHT data, surrogate variable analysis (SVA)^27^ identified a latent batch effect independent of the IR effect. This latent effect was included as a DESeq2 design term or corrected for data visualisation using the Limma package^28^.

### Proteomics quantification and differential abundance analysis

Proteomics analysis was performed by the NNRCO using LC-MS/MS to quantify protein abundance. Approximately 100 µg of proteins was normalized to 100 µl with the addition of 50mM ammonium bicarbonate followed by the addition of horse myoglobin standard (10 pmol) and 50 µl of 0.2% RapiGest were added. Samples underwent a reduction process with 10 mM dithiothreitol, followed by alkylation with 15 mM iodoacetamide for 30 minutes. Subsequently, digestion was carried out using sequencing grade modified trypsin (enzyme/substrate ratio: 1:50) at 37°C for 14-16 hours, quenched with 0.5% trifluoroacetic acid (45 minutes at 37°C), with the supernatant collected and dried. The dried peptides were reconstituted with 100 µl of 0.1% FA (1 µg/ µl), and 500 ng was loaded onto the column for liquid chromatography with tandem mass spectrometry (LC-MS/MS) analysis. Peptides were identified with PEAKS DB software^29^, and protein abundance was estimated from the summed AUC of identified peptides. We employed the DEP^30^ and proDA ^31^ R packages for further analysis. The DEP package filtered proteins with multiple missing values at default stringency, normalised abundance using VST, and imputed missing values for principal component analysis. The proDA package, which doesn’t require imputation for missing values, was used for all subsequent visualisation and differential abundance analysis. In the proDA workflow, raw abundance data were log-scaled and median normalised. This package utilises a probabilistic dropout model to include missing values in the analysis, which is conducted using linear modelling with empirical Bayesian priors to enhance the power of detecting differential abundance.

### Pathway-based analysis

Gene set enrichment analysis was conducted using the FGSEA package^32^, with genes ranked by the test statistic from differential expression analysis. Unlike typical analyses, lowly expressed genes were not removed to estimate NULL distributions in gene rankings accurately. For proteomics data, UniProt identifiers were first mapped to gene symbols. Pathways from Wikipathways, KEGG, and hallmark pathways were sourced from the Molecular Signatures Database^33^ and merged. The collapsePathways function in FGSEA was utilised to reduce redundancy by collapsing overlapping pathways. P-values were corrected using the Benjamini and Hochberg method, with FDR < 0.05 set as the significance threshold unless otherwise specified.

A single sample enrichment analysis was executed using the GSVA algorithm to highlight significant pathways visually. Genes contributing to enrichment (leading edge) were extracted and plotted as a heatmap with hierarchical clustering to represent each pathway’s impact at the gene level visually. The ComplexHeatmap^34^ package was used for this visualisation, employing default hierarchal clustering settings. Over-representation and protein-protein interaction (PPI) analysis were conducted using the online tool Metascape^35^ with default settings, with minor visual adjustments to PPI networks, such as font size and colour, made using Cytoscape^36^.

### Measurement of substrate metabolism

Glycolytic rates were measured by culturing cells or EHT with ^3^H-glucose (0.2 μCi/mL of D-[5-^3^H(N)] glucose) for 6 hours. Fatty acid oxidation rates were measured by culturing cells or EHT with ^3^H-oleate (0.2 μCi/mL of [9,10-^3^H(N)]-oleic acid) for 8 hours. Metabolic flux rates were measured by the rate of conversion of the ^3^H-substrate into ^3^H_2_O, as previously described^14^, using anion exchange chromatography for glycolytic measurements and Folch extraction for fatty acid oxidation measurements. Oil red O staining was carried out on EHTs using an Oil Red O staining kit (Sigma) after fixing in 4% PFA (nuclei were stained with DAPI).

### Western blotting

Cells or EHT were washed with DPBS and cultured for 30 mins with or without 10 μg/ml of insulin, followed by lysing and snap freezing in lysis buffer. Western blotting was carried out to measure pAkt (Abcam Ab81283) and total Akt (Abcam Ab32505) and expressed as the ratio of pAkt/Akt.

### Measurement of EHT contractile function

Videos of contracting EHT were taken at 60x magnification for 15-30 seconds using a Canon EOS 100D camera. The videos were then converted into 50 frames-per-second tiff sequence images using a terminal in MacOS. They were then processed using Musclemotion software v.1195–197 as a macro plug-in FIJI/ImageJ software^37^. The data generated was used to calculate contraction duration and relaxation time.

### Live/dead staining of cells

To determine if inducing IR was causing cell death, we used a live/dead staining procedure that was carried out using Readyprobes® (ThermoFisher Scientific), according to the manufacturer’s protocol. Readyprobes® consists of NucBlue® Live reagent and NucGreen® Dead reagent, and labelled cells were imaged using fluorescence microscopy using the DAPI and GFP filters, respectively.

### Statistical analysis and plots

All statistical analysis (except over-representation) was conducted in R (v 4.3.1). Statistical comparisons displayed in boxplots for genes demonstrate DESEQ2 results and other comparisons were made using the Wilcoxon Rank-Sum method. Boxplots depict median and interquartile ranges. Barplots depict means with the standard deviation, with student’s t-test comparisons made using the ‘stat_compare_means’ function from ggpubr. The levels of significance denoted by asterisks are as follows: < 0.0001 ****, < 0.001 ***, < 0.01 **, < 0.05 *, > 0.05 ns. Additional plots were generated using complexHeatmap and EnhancedVolcano packages. Illustrations were created with BioRender.com.

## Results

### Culturing 2D hiPSC-CM in an “insulin-resistance” media captures many characteristics of the DbCM phenotype

During the pathological progression of type II diabetes mellitus (T2DM), the heart becomes exposed to hyperglycaemic, hyperlipidaemic, and hyperinsulinaemic environments, which dysregulates cardiac metabolism and contractile function leading to diabetic cardiomyopathy (DbCM)^4^. Therefore, we utilised human induced pluripotent stem cell-derived cardiomyocytes (hiPSC-CMs) exposed to an ‘insulin resistance’ (IR) media that recapitulates these systemic factors (Figure 1A) to identify human relevant mechanisms related to diabetic cardiomyopathy (DbCM) *in vitro*. We used transcriptomics to undertake a high-dimensional assessment of our model and determine the broad changes underlying the disease pathology. Principal component analysis established a clear separation of samples by control or IR media treatment, with the effect captured across the main component and explaining 59% of the variation (Figure 1B). Differential expression analysis identified 2121 significantly downregulated and 2021 significantly upregulated genes (DEGs) in the IR group compared with the control group (Figure 1C). Given that 18854 genes were tested, these significant changes represent ∼23% of all non-lowly expressed genes being altered. Therefore, the transcriptomics analysis captures global differences in gene expression induced by the IR media in the hiPSC-CM model.

**Figure 1.**
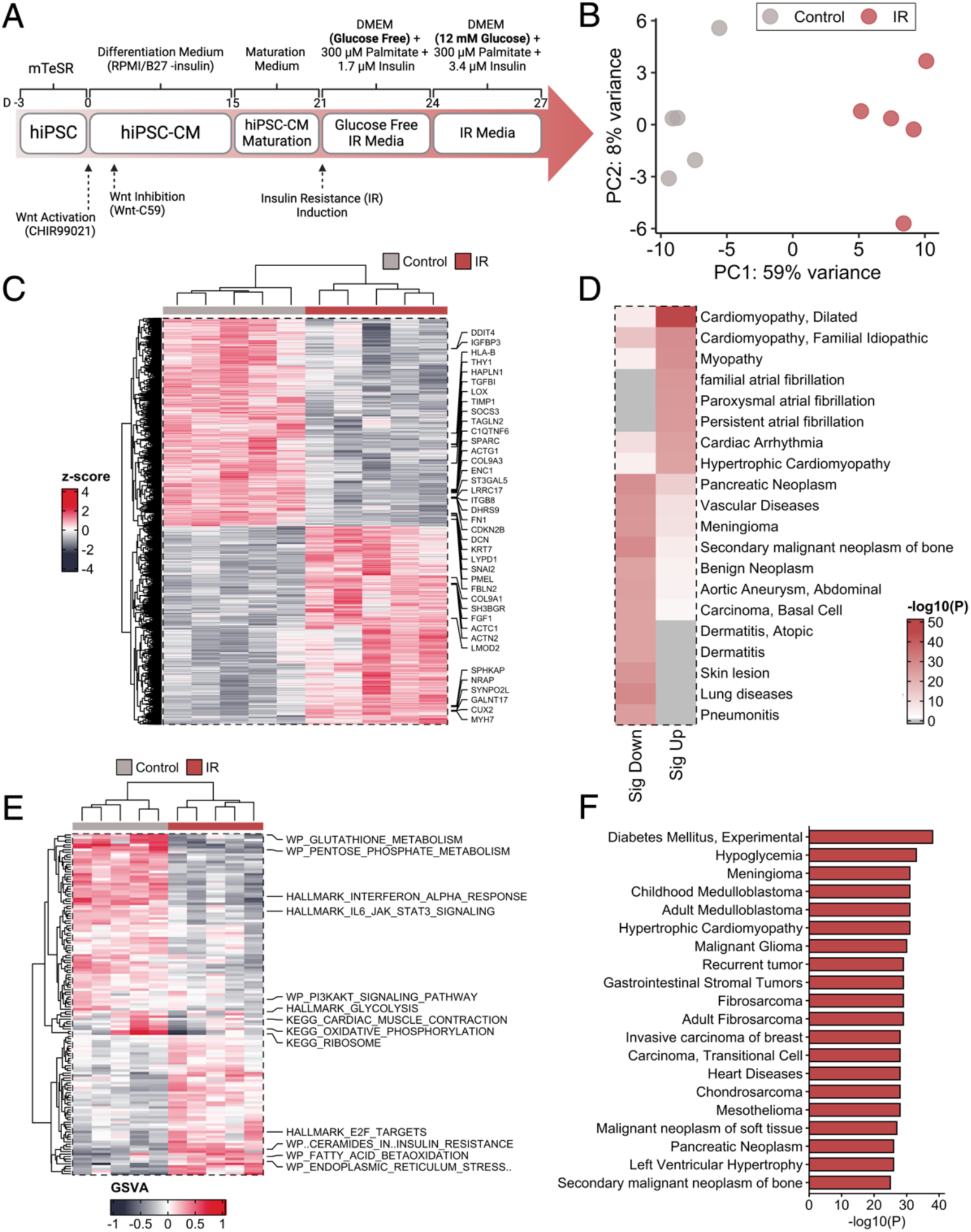
Culturing 2D hiPSC-CM in an IR media captures many characteristics of the DbCM phenotype. Insulin-resistant hiPSC-CMs were generated over a fifteen-day differentiation, seven-day maturation and seven-day insulin resistance protocol (A). Principal component analysis from transcriptomics of insulin-resistant and control hiPSC-CMs shows separation of the groups across the first component (B). Differential expression analysis established genes significantly altered by insulin resistance (C). Cardiomyopathy is significantly enriched for disease-related pathways overrepresented in significantly different genes (D). Single sample summary scores for non-overlapping pathways enriched in all genes ranked by their differential expression separated the groups and included DbCM-related terms (E). Diabetes and DbCM-related disease terms were most significantly overrepresented in the leading-edge genes for select pathways defined in the main text (F).

Given the large-scale transcriptional changes observed, a pathway approach was utilised to reduce complexity and contextualise these changes to broader biological functions. Several cardiovascular disease-related terms were significantly over-represented within the differentially expressed genes, such as those relating to cardiomyopathies (Figure 1D). We continued this approach by establishing the pathways enriched within the top or bottom of the complete gene list ranked by their differential expression between control and IR. This methodology is advantageous as it considers all expressed genes and can detect pathways changing in coordinated but small ways that would be otherwise missed. One hundred and forty-seven pathways were significantly enriched after correcting for multiple tests and collapsing overlapping terms. Single sample enrichment analysis allowed the expression of each pathway to be visualised and identified clear separation across the treatment groups with hierarchical clustering (Figure 1E). Multiple metabolic, signalling, and biological process terms related to the development of T2DM were identified. Interestingly, the genes contributing to the enrichment (leading-edge) for glycolysis, oxidative phosphorylation, β-oxidation, endoplasmic stress, PI3K-Akt and insulin signalling pathways were collectively over-represented for ‘Diabetes Mellitus, Experimental’ in the DisGeNET gene sets (Figure 1F). This IR protocol did not induce cell death or apoptosis in the human iPSC-CM (Supplementary Figure 1). Taken together, the IR 2D hiPSC-CM model captures many characteristics of the DbCM phenotype.

### The 2D hiPSC-CM model captures the IR phenotype with decreased glycolysis

Insulin signalling is mediated by the PI3K-Akt pathway, and IR is a feature of the diabetic heart. Therefore, we investigated whether genes downstream of insulin signalling were differentially regulated at the transcriptional level in our model. The expression of the leading-edge genes for the PI3K-Akt pathway was visualised as this gene set displayed significant negative enrichment in the IR model (Figure 2A, B). Hierarchical clustering for their expression separated the treatment groups and was notably decreased within the IR media group. To validate these gene findings and provide functional confirmation of IR in this model, the pAkt/Akt ratio was measured via western blotting following insulin stimulation. It was significantly decreased by 29% in the IR cells compared with control cells (Figure 2C).

**Figure 2.**
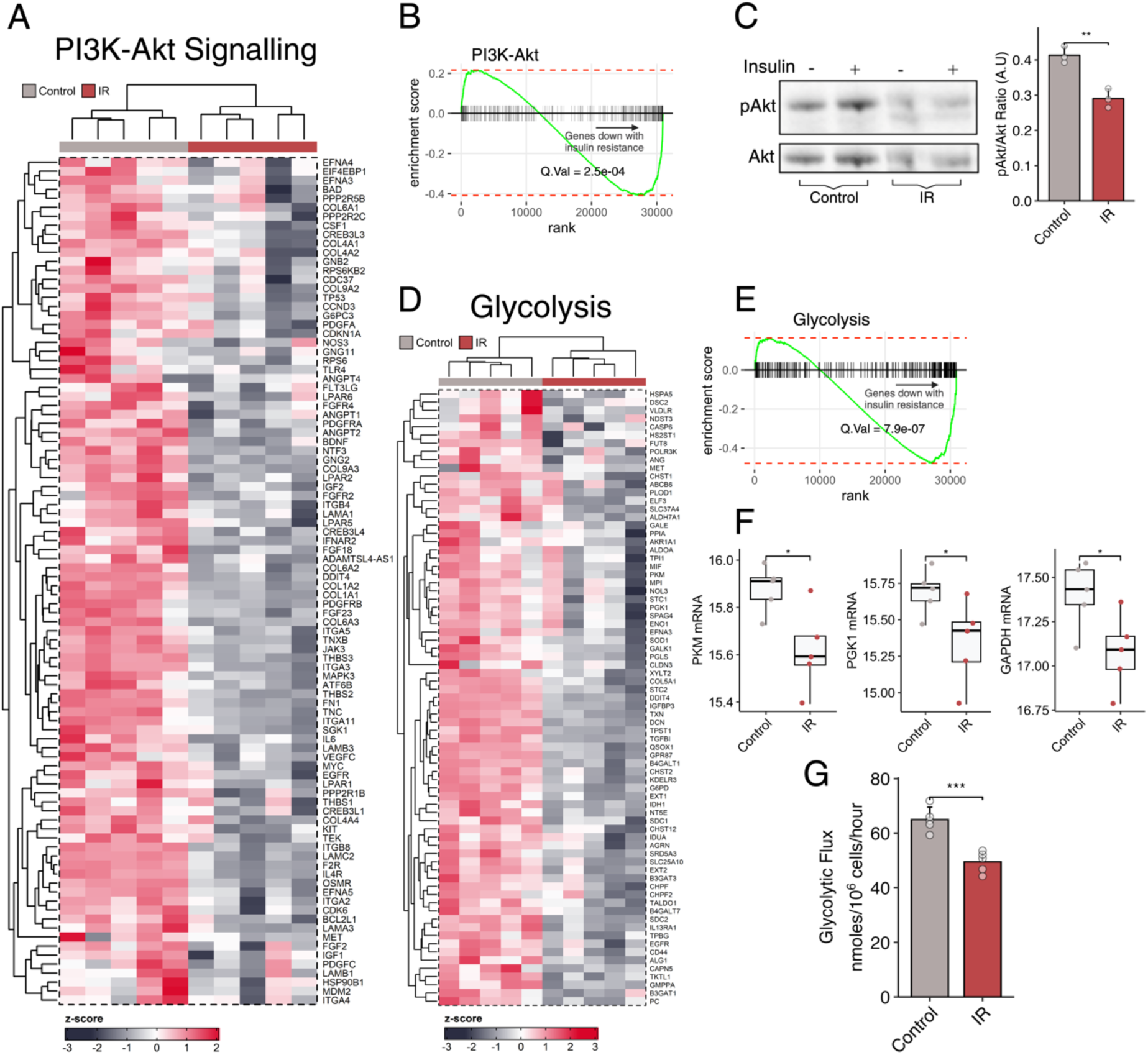
The 2D hiPSC-CM model captures the IR phenotype demonstrating decreased glycolysis. The leading-edge genes for the significantly negatively enriched PI3K-Akt pathway are visibly reduced in the insulin-resistant samples (A). The enrichment plot visualises the enrichment of PI3K-Akt genes at the bottom of the gene list (B). Western blots show the level of phosphorylated Akt after insulin exposure was significantly blunted for insulin-resistant hiPSC-CMs (C). mRNA expression of the glycolysis pathway was significantly blunted in the insulin-resistant hiPSC-CMs, as shown on a pathway (D, E) and rate-limiting enzyme level for pyruvate kinase M (PKM), phosphoglycerate kinase 1 (PGK1) and glyceraldehyde-3-phosphate dehydrogenase (GAPDH) (F). Radioisotope flux measurements confirm significantly blunted glycolysis in insulin-resistant hiPSC-CMs (G).

Insulin regulates glucose metabolism in the heart, and patients with T2DM have decreased glycolysis. In our IR media hiPSC-CM model, the glycolysis pathway showed significant negative enrichment and notably reduced expression in the IR media-treated samples compared with the control samples (Figure 2D, E). In agreement, several glycolytic enzymes were significantly decreased in the IR group (Figure 2F). Glycolytic flux was measured using radioisotope tracers to validate the transcriptomics data, and glycolytic rates were significantly reduced by 23% in the IR group compared with the control group (Figure 2G).

### The 2D IR model demonstrates a metabolic shift towards increased fatty acid utilisation and changes in mitochondrial oxidative phosphorylation

In contrast to glycolysis, expression of the fatty acid metabolism β-oxidation pathway was positively enriched within our IR model, showing an increased expression that separated the treatment groups robustly with clustering (Figure 3A, B). The leading-edge genes included several rate-limiting enzymes, significantly increasing within our model (Figure 3C). Aligning with these findings, the PPAR signalling pathway showed significant positive enrichment (FDR = 0.03; NES = 1.6), but was removed whilst collapsing overlapping pathways. Therefore, to ensure a non-redundancy, single sample enrichment evaluated all known gene targets for the master regulator of fatty acid metabolism, PPARα. These genes collectively increased significantly within the IR model (Figure 3D), with an example of the known downstream target of PPARα signalling, PDK4, shown. Fatty acid oxidation rates were measured using radioisotope tracers to validate the transcriptomics data, and fatty acid oxidation rates were significantly increased in the IR group by 2.9-fold compared with the control group (Figure 3E). In patients with DbCM, defects in myocardial energetics have been identified and associated with mitochondrial dysfunction. Despite significant negative enrichment for the oxidative phosphorylation pathway in our IR model, the expression for these genes did not robustly cluster the treatment groups (Figure 3F, G). However, several electron transport chain subunits demonstrated significantly altered expression levels (Figure 3H). Among those significantly changing, all but the succinate dehydrogenase enzyme complex (SDHA) and mitochondrially encoded NADH dehydrogenase 1 (MT-ND1) were negatively expressed in the IR model.

**Figure 3.**
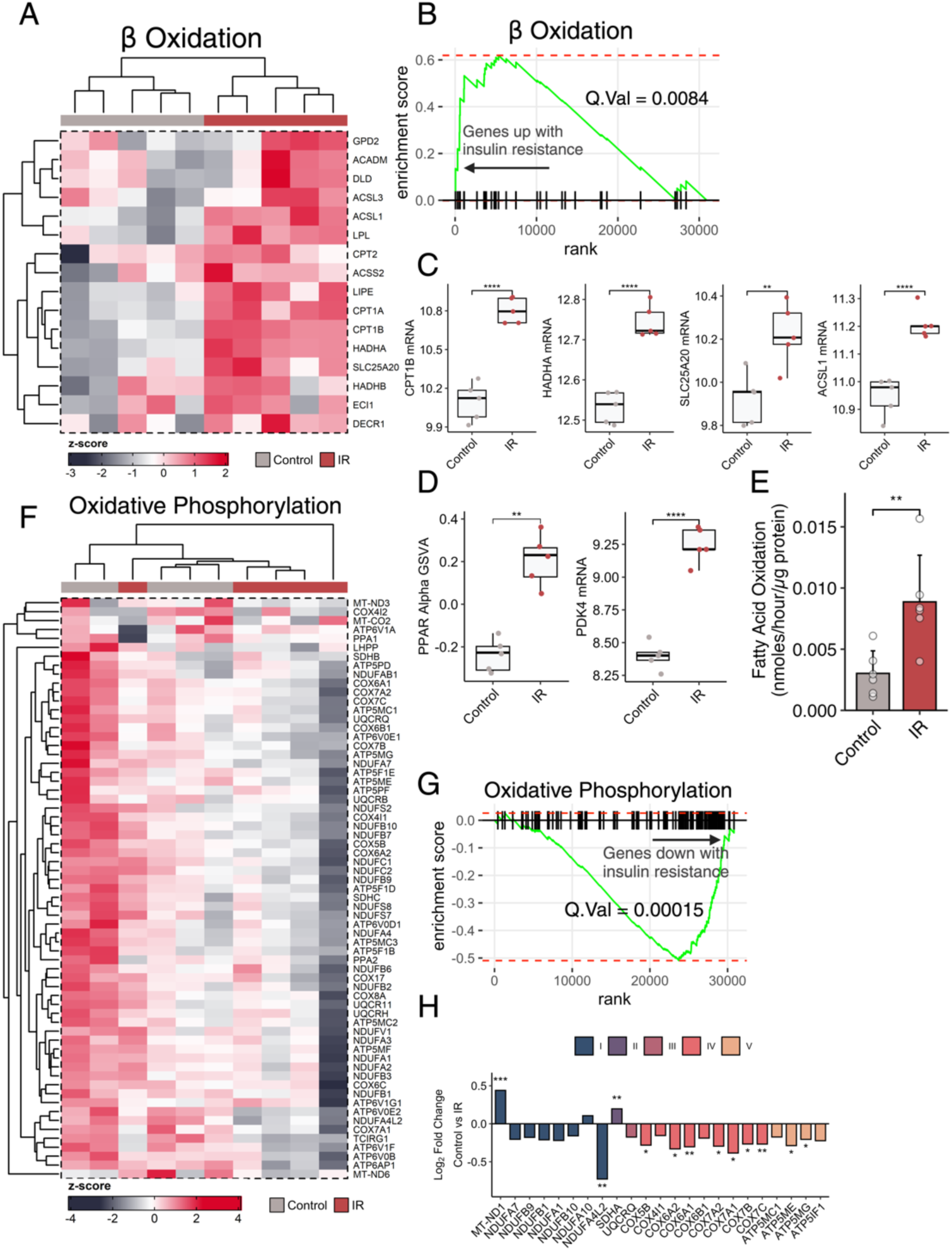
The 2D IR model demonstrates a metabolic shift towards increased fatty acid utilisation and changes in mitochondrial oxidative phosphorylation. Gene expression for the fatty acid oxidation pathway was significantly induced by insulin resistance on a pathway (A, B) and rate-limiting enzyme level for carnitine palmitoyl transferase 1B (CPT1B), hydroxyacyl-CoA dehydrogenase A (HADHA), carnitine/acylcarnitine translocase (SLC25A20) and long-chain acyl-CoA synthetase 1 (ACSL1) (C). Gene targets for the fatty acid regulatory transcription factor PPARα, and one of its downstream targets, pyruvate dehydrogenase kinase 4 (PDK4), were significantly increased with insulin resistance (D). Radioisotope flux measurements of fatty acid oxidation were also significantly elevated (E). Despite significant negative enrichment for the oxidative phosphorylation pathway, the expression for leading-edge genes did not robustly separate the groups via hierarchical clustering (F, G). Most mitochondrial genes showed significantly reduced expression with insulin resistance, except for SDHA and MT-ND1, which were induced (H).

Using MUSCLEMOTION software, we assessed contractile function in the 2D hiPSC-CMs cultured in control and IR media (Supplementary Figure 2). There were no significant differences in contraction duration or relaxation time, but there was a lot of variation in the data and heterogeneity across the plate.

### Culturing 2D hiPSC-CMs in IR media blunts the response to hypoxia

Patients with T2DM have a more rapid progression into heart failure after myocardial infarction (MI)^38^. Activation of the hypoxic response is critical for remodelling post-MI^39^, and this process is blunted in T2DM^40,41^. Therefore, the cellular transcriptional adaptation to hypoxia was investigated in the control and IR hiPSC-CMs. PCA demonstrates that the transcriptional response to hypoxia was captured and represents a more pronounced treatment effect, explaining 62% of the variation across the first component compared to the 15% in component two, separated by IR media (Figure 4A). Interestingly, the degree of separation across component two is more substantial for the hypoxic samples than normoxic samples, suggesting an interaction effect with hypoxic-specific changes occurring within the IR model. Furthermore, the hierarchical clustering of sample distances suggests that the hypoxic IR samples relate more closely to the normoxic groups than the hypoxic control samples, indicating an altered response to hypoxia in the IR cells (Figure 4B).

**Figure 4.**
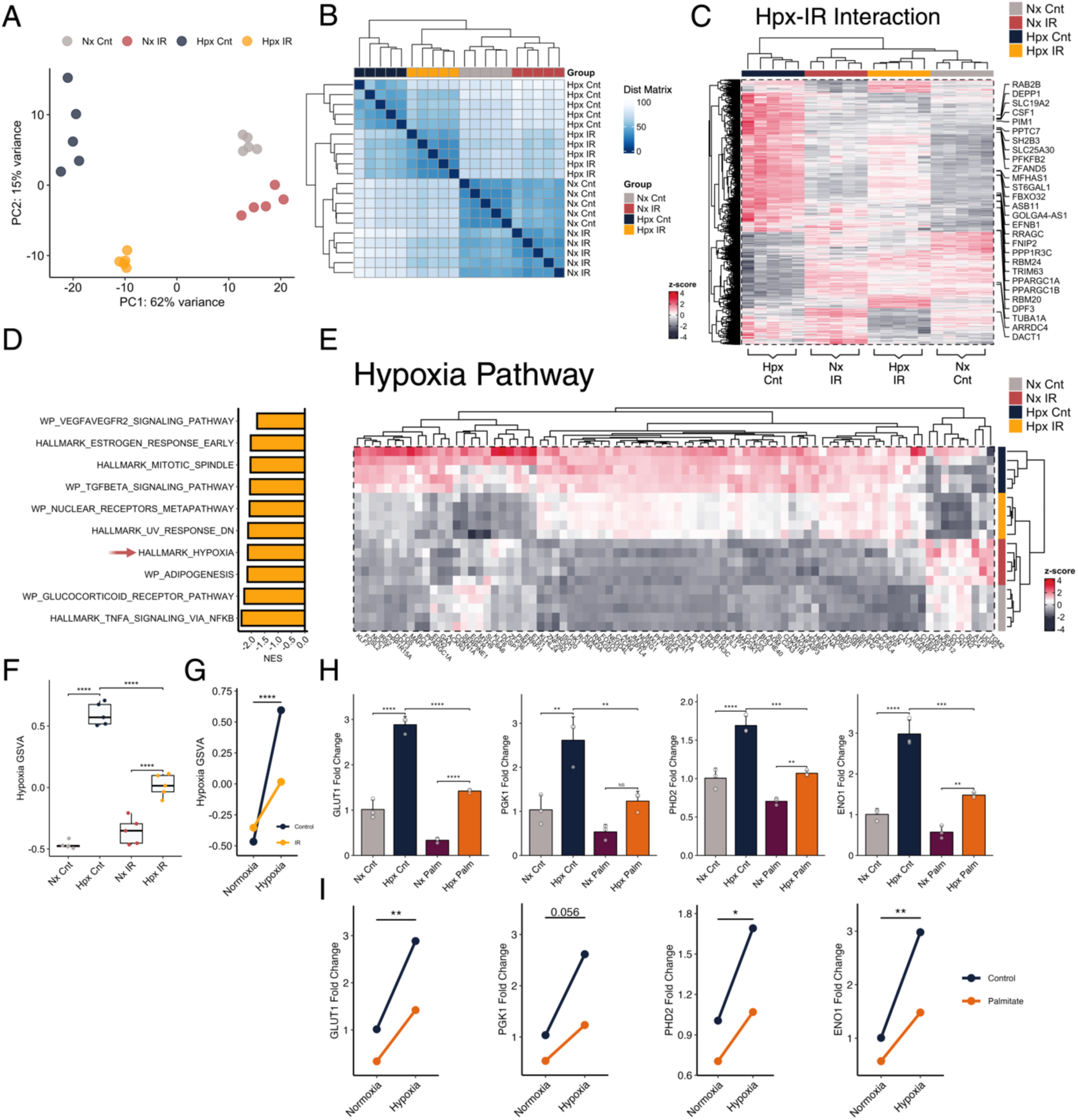
Culturing 2D hiPSC-CMs in IR media blunts the response to hypoxia. Principal component analysis of transcriptomics for control and insulin-resistant hiPSC-CMs in normoxia and hypoxia (A). In the hierarchical clustering of sample distances (B), the insulin-resistant hypoxia group clusters closer to the normoxic groups than the hypoxic control group. Differential expression analysis identifies the genes with significant interaction effects for insulin resistance and oxygen concentration (C). Hypoxia-related terms are significantly enriched in genes ranked by their interaction effect (D). Visualising the hypoxia pathway demonstrates the blunting effect of hypoxic induction with insulin resistance across the leading-edge genes (E) and summary scores for the pathway (F, G). Culturing in increased concentrations of palmitate alone can replicate this blunting effect seen in IR hiPSC-CMs, as demonstrated using qPCR of hypoxia-induced genes, including glucose transporter 1 (GLUT1), phosphoglycerate kinase 1 (PGK1), prolyl hydroxylase domain 1 (PHD1) and enolase 1 (ENO1) (H, I).

To further examine the relationship between hypoxia signalling and IR, differential expression analysis was undertaken with an interaction term to identify genes altered by the IR phenotype in a hypoxia-specific manner. Strikingly, 3034 genes differed significantly, and visualising their expression revealed that culturing in IR media blunted the induction for most of these genes in hypoxia (Figure 4C). The downstream enrichment analysis showed that hypoxia and several related pathways were significantly enriched among the genes affected by this interaction (Figure 4D). Again, visualising the genes within the hypoxia pathway identified a blunting of hypoxia signalling in the IR cells (Figure 4E). In support, the hierarchical clustering of samples by their expression for these hypoxia-related genes recapitulated the dendrogram reported earlier (Figure 4B), with IR hypoxic samples clustering closer to normoxic groups. GSVA summary scores for these genes demonstrate that both control and IR hiPSC-CMs had significant induction of the hypoxic response (Figure 4F). However, the hypoxic IR group was significantly blunted compared with hypoxic control. The interaction plot further visualises this effect (Figure 4G).

The protocol for inducing the IR phenotype within hiPSC-CMs utilises high concentrations of insulin, glucose, and fatty acids. However, culturing with palmitate alone was sufficient to recapitulate the blunting of hypoxia signalling in hiPSC-CMs. Utilising qPCR, several known hypoxia-inducible factor (HIF) targets showed a significant reduction in hypoxic induction in hiPSC-CM cultured with palmitate (Figure 4H). Furthermore, significant interactions between oxygen and media conditions were identified for most targets (Figure 4I). Therefore, these results validate our sequencing findings and suggest the blunting of the response to hypoxia observed in our IR model is primarily driven by exposure to high palmitate concentrations, akin to dyslipidaemia in T2DM patients.

### Integrated transcriptomics and proteomics demonstrate network hubs in 3D engineered heart tissue (EHT) that capture the disease phenotype

Culturing hiPSC-CMs as 3D-engineered heart tissue (EHT) replicates the increased workload and shear stress of *in situ* cardiomyocytes by providing restraining posts between which the tissue contracts, which confers increased maturity (Figure 5A). Thus, an IR model utilising EHTs may have improved translation to the pathology in T2DM patients. However, this may be outweighed by increased cost, time, and potential for introducing variability. Therefore, we developed an IR EHT model, utilising the same IR media and exposure protocol developed in the monolayer, and assessed the model using parallel transcriptomics and proteomics.

**Figure 5.**
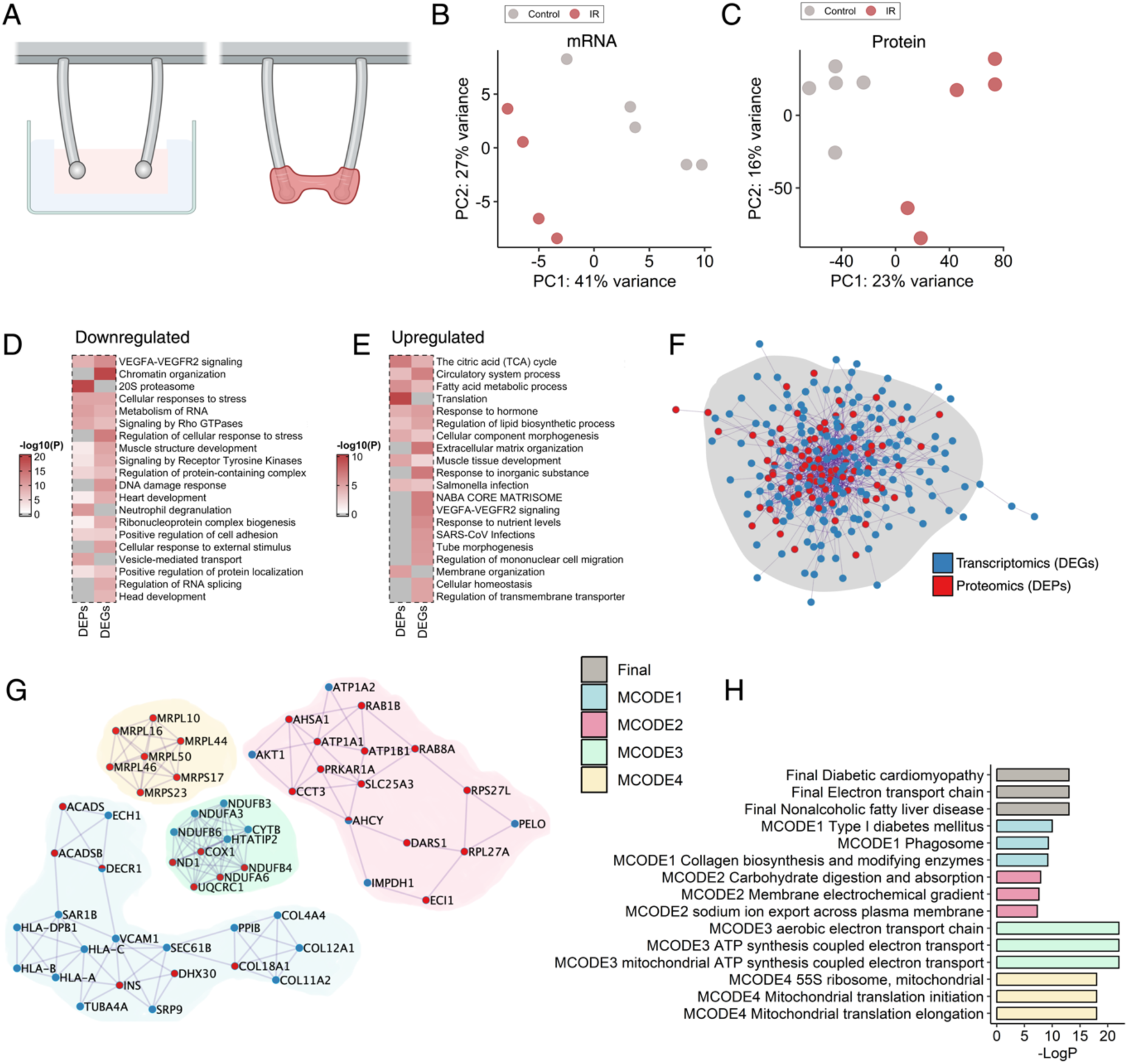
A 3D Engineered Heart Tissue (EHT) model of insulin resistance captures the disease phenotype, demonstrating integrated networks of genes and proteins related to diabetic cardiomyopathy. EHTs were generated using coagulation factors to solidify hiPSC-CMs within agarose moulds (A). Principal component analysis for transcriptomics (B) and proteomics (C) captures the insulin-resistance eZect across the main component. Overrepresented pathways within negatively diZerentially expressed genes (DEGs) or proteins (DEPs) (D) or positive DEGs or DEPs (E) capture several metabolic-related terms. Based on known protein-protein interactions, an integrated network of upregulated genes (protein coding) and proteins was constructed (F). Dense subnetworks within the final network were established with the MCODE algorithm (G). Cardiomyopathy and metabolic-related pathways were significantly overrepresented within these networks (H).

Principal component analysis of normalised transcriptomics and proteomics identified a separation of control and IR EHT samples across the main component (Figure 5B, C). Differential abundance and expression analysis identified the proteins and genes changing between the control and IR groups. For transcriptomics, 321 genes were significantly downregulated, and 224 genes were significantly upregulated by IR media. Meanwhile, only eleven proteins significantly differed between control and IR EHT when corrected for multiple comparisons by controlling for the false discovery rate (FDR). Therefore, the 132 downregulated and 109 upregulated proteins (DEPs) at the raw p-value threshold were used in the downstream analysis with the limitation that FDR was uncontrolled. Multiple pathways were significantly over-represented within the downregulated genes or proteins (Figure 5D) and upregulated genes or proteins (Figure 5E). Of note, terms for signalling by receptor tyrosine kinases (the family for the insulin receptor) and vesicle-mediated transport (involved in glucose uptake) were overrepresented in the downregulated sets. Fatty acid metabolism and extracellular matrix remodelling terms were overrepresented in the upregulated sets. A protein-protein interaction approach improved the power of over-representation analysis by integrating the upregulated DEGs and DEPs into an overall network (Figure 5F) and dense subnetworks (Figure 5G). Within the broad network, the diabetic cardiomyopathy pathway was significantly overrepresented; several metabolic and diabetes-related pathways were overrepresented within the subnetworks (Figure 5H). Overall, initial pathway analysis suggests the 3D IR EHT model has also captured the general phenotype of the T2DM heart.

### The IR EHT model demonstrates a metabolic shift from glucose to fatty acids accompanied by diastolic dysfunction

Aligning with our findings in the monolayer, the insulin signalling pathway demonstrated significant negative enrichment in the IR 3D EHT model (FDR = 0.037; NES = -1.49) and robustly clustered the samples by treatment (Figure 6A). Confirming these expression changes related to a repression of insulin signalling, phosphorylated Akt to Akt ratios were reduced by 80% following insulin stimulation in the IR EHT (Figure 6B, C). Furthermore, in line with the physiological role of insulin in regulating glucose metabolism, glycolytic flux rates were decreased by 46% in the IR EHTs compared with control EHTs (Figure 6D). The metabolic shift towards increased fatty acid metabolism was also replicated for our EHT model. The fatty acid metabolism pathway showed positive enrichment in the IR EHTs at the mRNA (FDR = 0.00093; NES = 1.78) and protein levels (P value = 0.0093; NES = 1.75), robustly clustering the treatment groups in both (Figure E, F). These protein and gene expression changes culminated in increased rates of fatty acid oxidation in the IR EHT by 32% compared with control EHT (Figure 6G). Intramyocardial triglyceride stores are increased in patients with T2DM and have been associated with a lipotoxic phenotype that negatively impacts diastolic function. The IR EHT oil red O staining demonstrated accumulation of intracellular lipids, with a 2-fold increase in staining intensity compared with controls (Figure 6H, I). The PPAR signalling pathway, which regulates fatty acid metabolism, showed significant positive enrichment in the IR EHTs, as demonstrated for CD36, the PPARα target critical for fatty acid uptake (Figure 6J, K).

**Figure 6.**
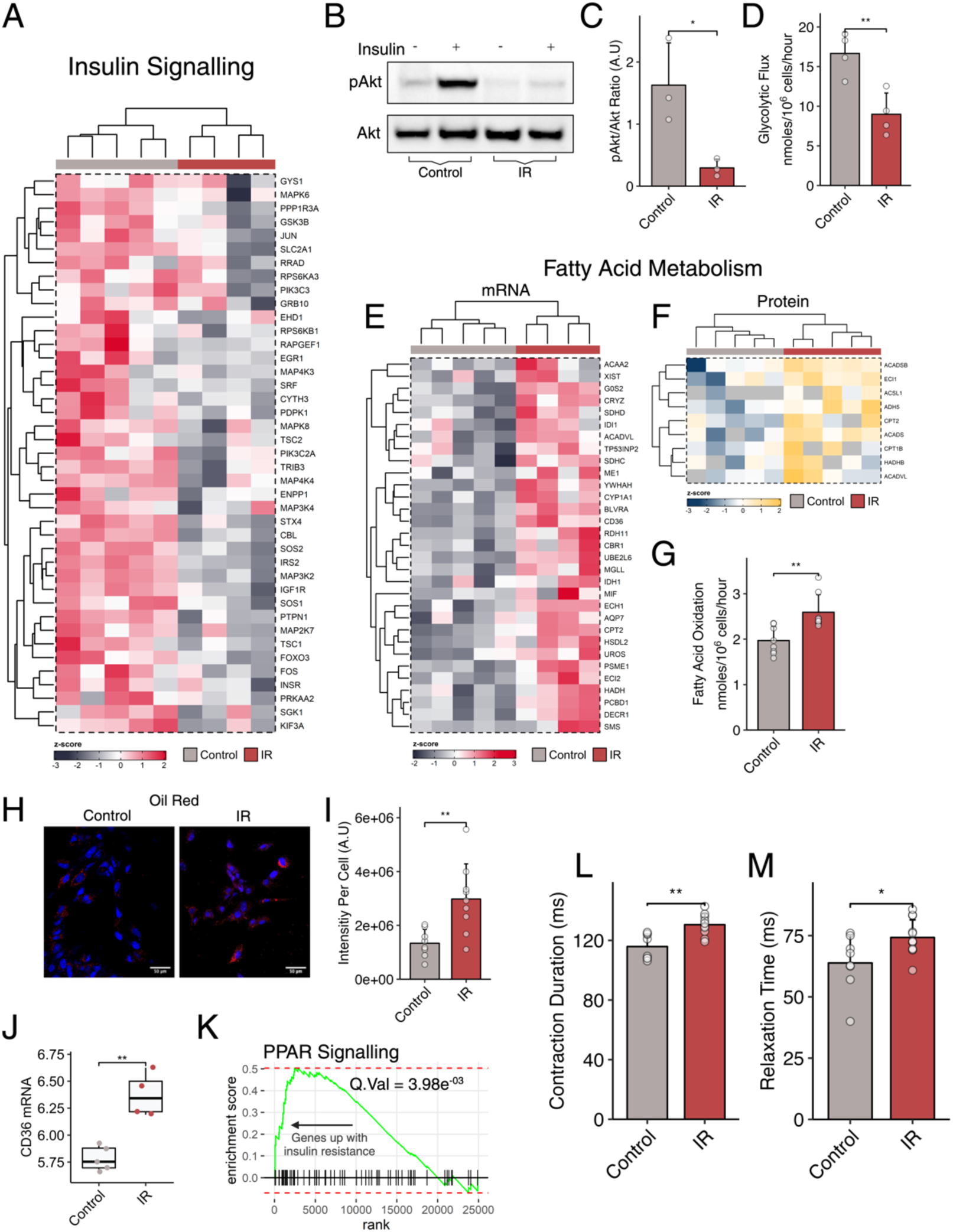
The IR EHT model demonstrates a metabolic shift from glucose to fatty acids accompanied by diastolic dysfunction. Expression for the leading-edge insulin signalling pathway genes visualises the negative enrichment of this pathway for insulin-resistant EHTs (A). Protein levels for the pAkt/Akt ratio were significantly lower for insulin-resistant EHTs than controls after insulin stimulation (B, C). Glycolytic flux, as measured with radioisotopes, was also significantly reduced (D). Insulin-resistant EHTs demonstrated induction for fatty acid metabolism on a transcriptomic (E) and protein level (F). Radioisotope measurements of fatty acid oxidation were increased in insulin-resistant EHTs (G). Oil red o staining identified significantly increased lipid droplet accumulation in insulin-resistant EHTs (H, I). Mediators of lipid uptake and fatty acid metabolism, CD36 and PPAR signalling were significantly increased with insulin resistance (J, K). Musclemotion captured measures of contraction duration (L) and relaxation time (M), which significantly increased for insulin-resistant EHTs.

Diastolic dysfunction is a known hallmark of DbCM, and a vital advantage of a 3D model is the ability to take more physiologically relevant measures of contractile function *in vitro*. In the 3D IR EHT, contraction duration and relaxation time increased by 13% and 16%, respectively, compared with control EHT (Figure 6L, M).

## Discussion

Human-centric models of DbCM are needed for mechanistic studies of disease development and testing of new therapies. Here, we have developed a model of DbCM by culturing hiPSC-CM in 2D or 3D in an “insulin resistance” (IR) media, and used a systems biology approach to take a broad view of the regulated pathways affected. We demonstrate that culturing hiPSC-CM in our IR media activates many of the pathways implicated in DbCM, including a metabolic shift from glucose towards fatty acid metabolism, mitochondrial dysfunction, extracellular matrix remodelling, ER stress, a blunted response to hypoxia and diastolic dysfunction.

### Culturing cells in IR media induces a transcriptional profile identified as Diabetes Mellitus

hiPSC-CMs have traditionally been used to study monogenic diseases by generating CM directly from affected patients or introducing known mutations into healthy cells. Using hiPSC-CM to model systemic complex multifactorial diseases has received much less attention. By mimicking the hyperglycaemia, hyperlipidaemia and hyperinsulinemia present in patients with T2DM, we hoped to impact a range of different cellular processes affected within the heart in T2DM, taking a broader rather than a narrower approach to mimicking this multifactorial disease. However, we acknowledge that there are other circulating factors we haven’t accounted for in our model, including adipokines, cytokines and inflammatory cells. Using an unbiased multi-omics approach, we could take a comprehensive view of pathways modulated by our IR protocol and investigate if they mirror the cardiac diabetic phenotype seen in human and animal models. Notably, multiple metabolic, signalling, and biological process terms related to the development of T2DM were identified, resulting in ‘Diabetes Mellitus, Experimental’ being the primary disease identifier in the DisGeNET gene sets.

### 2D and 3D IR hiPSC-CMs demonstrate the metabolic shift from glucose to fatty acid utilisation

One of the hallmarks of the heart in T2DM is a metabolic shift away from glucose use towards an increased dependence on fatty acid metabolism^5,42^. This metabolic shift occurs early during disease progression^5^. It has been shown to contribute to diastolic dysfunction, with preclinical studies to rebalance metabolism correcting diastolic function^43,44^. In the 2D and 3D IR models, the shift from glucose towards fat metabolism was evident, with transcriptomic changes reflecting decreased glycolytic flux and increased fatty acid oxidation flux. The increase in fatty acid metabolism was evident in both the transcriptomic and proteomic profiling, likely driven by increased PPARα activation, a key driver of the diabetic metabolic phenotype^45^. Regulation of lipid biosynthetic processes was increased in the DEPs and DEGs, which contribute to the increased oil red O staining for triglycerides and upregulation of pathways linking ceramides with IR. Critically, the IR media developed in this study induced IR in both 2D and 3D EHT cultures, as evidenced by the downregulation of insulin signalling cascade genes and proteins, and validated by decreased insulin-induced phosphorylation of Akt. Thus, the metabolic immaturity of the 2D versus 3D hiPSC did not influence the use of this model in recapitulating the metabolic phenotype of DbCM.

Culturing cells in high concentrations of fatty acids induces apoptosis and cell death^46^. Therefore, various iterations of the protocol were developed to choose the one that induced IR without causing cell death and apoptosis. One important factor is that our cells mature for 7 days in a low fatty acid concentration (80µM) to replicate the foetal to adult metabolic shift towards oxygen-dependent metabolism. Therefore, they have already upregulated the fatty acid metabolic and mitochondrial respiratory pathways before assigning them to control or IR treatment groups. This means the cells are already primed and able to metabolise fats, and we suggest this causes less “fat shock” to the cells, resulting in minimal induction of apoptosis and death in the IR group.

### The IR model impacts key mediators implicated in the development of DbCM

DbCM is characterised by diastolic dysfunction, hypertrophy, and diffuse fibrosis, with critical mediators of DbCM including abnormal metabolism, endoplasmic reticulum stress, extracellular matrix (ECM) remodelling, dysregulated calcium handling and oxidative damage^4^. In our IR model, several of these mediators were identified in our omics-based profiling. Genes involved in ER stress and ECM remodelling were upregulated, whereas genes involved in oxidative defence mechanisms were downregulated. Notably, in the 3D EHT, both proteins and genes involved in ECM remodelling and collagen biosynthesis were upregulated, forming a dense subnetwork. In diabetes, diffuse fibrosis is associated with the activation of fibroblasts and collagen synthesis^47^. Despite efficient differentiation, some non-myocytes remain within differentiated cell populations and those that remain after glucose depletion may increase by proliferation^48,49^. Thus, it is highly likely that our culture is not exclusively hiPSC-CMs but also contains hiPSC-fibroblasts. However, this is a serendipitous advantage for the study of IR, given the critical role of fibrosis in the developing DbCM disease.

### Blunted response to hypoxia in IR recapitulates impaired post-MI remodelling seen in patients and animal models

Hypoxia is a critical component of ischaemia, and activation of the hypoxia-inducible factor occurs rapidly during an MI and is sustained through into the development of heart failure, driving hypoxic adaptation critical for post-MI remodelling^50^. In diabetic patients with ischaemia, activation of HIF1α is blunted^40^, and in animal models, the induction of the hypoxic response is suppressed in T2DM^51^. Hypoxia induced a greater transcriptional response in the hiPSC-CMs than that mediated by IR, indicating that our nutrient-overloading culture conditions were a relatively mild challenge for these cells. The data demonstrated a clear interaction between IR and hypoxia, with IR hiPSC-CMs having a much more stunted response to hypoxia than control hiPSC-CMs. Additionally, hierarchical clustering of sample distances demonstrated that exposing the IR hiPSC-CMs to hypoxia made them resemble the normoxic groups more closely than the hypoxic control hiPSC-CMs. This agrees with previous animal studies that have housed T2DM rats in chronic hypoxia^51^. In mouse cardiomyocytes, it has been shown that hyperlipidaemia drives the blunted response to hypoxia^41^. For several HIF targets, we demonstrate that hyperlipidaemia alone can blunt the induction of hypoxia genes in hiPSC-CMs, and explains why pharmacological fatty acid uptake inhibitors are beneficial for HIF activation in IR^41^. This blunted activation of HIF pathways in human cardiomyocytes provides further translational relevance for the beneficial effects of *in vivo* pharmacological HIF activators in the T2DM rat^17^.

### Advantages and disadvantages of modelling IR in 2D vs. 3D

The 3D EHT recapitulated many signalling pathways implicated in DbCM and demonstrated functional readouts of diastolic dysfunction, displaying increased relaxation time and contraction duration. Various preclinical studies developing novel therapeutic approaches to improve the heart in diabetes have shown that directly targeting and rebalancing metabolism improves diastolic function^43,44,52–54^. Thus, developing a human-centric IR hiPSC-CMs model that unifies abnormal metabolism and diastolic dysfunction is advantageous for drug development and testing. The 2D IR hiPSC-CMs model, while recapitulating the signalling pathways implicated in DbCM, did not demonstrate diastolic dysfunction, an apparent downside to using the 2D model if functional readouts are desirable. For the 2D cells, the limitation of the functional readouts was the heterogeneity of the data, as there was minimal synchronicity in contraction and relaxation across the plate (unlike when the cells were cultured in EHT). A potential downside of the IR EHT is that we observed fewer significantly differentially expressed genes, which may suggest greater heterogeneity for 3D vs. 2D. Despite EHTs being generated from the same differentiation batch^55^, we postulate that gradients within individual EHTs, the larger number of cells required to generate the EHT, or the process of dissociation and formation of the EHT may introduce increased variation. It is essential also to consider the extra cost and time needed to develop the 3D model from the 2D model, especially when an increased sample size may be required to compensate for increased variation. Our viewpoint is that 2D and 3D models have advantages over the other, and the better model depends on the scientific question the investigator addresses. In our opinion, the 2D IR model is better suited to addressing human-centric mechanisms of DbCM, whereas the 3D EHT IR model is better suited to drug testing and functional translation.

## Acknowledgements

We want to thank Jack Miller for the constructive discussion on statistics.

---

**Supplementary Figure 1.**
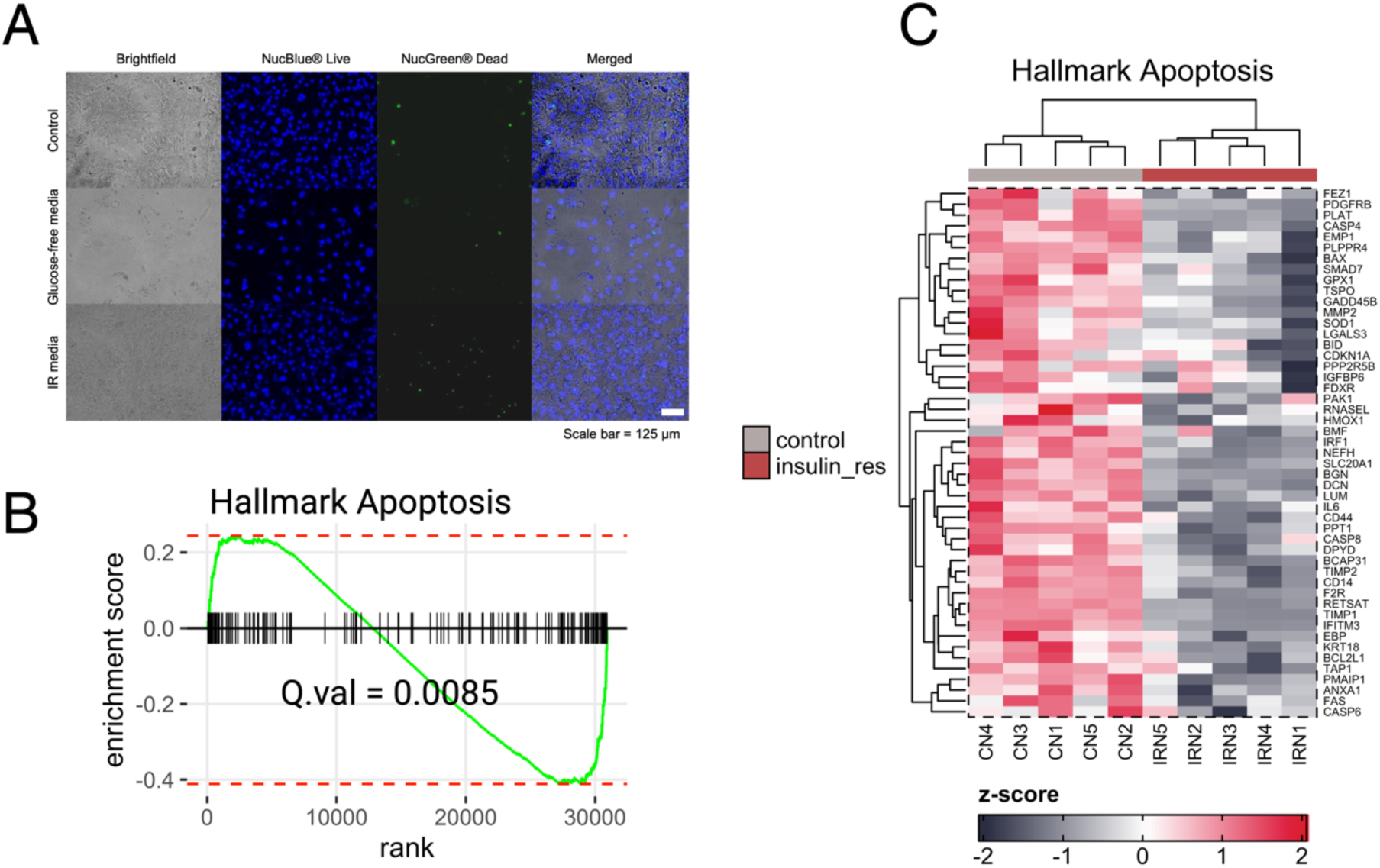
Cell death and apoptosis were not induced by culturing in IR media. Fluorescence microscopy of live/dead staining shows limited induction of cell death with the insulin-resistance protocol after 3 days (labelled Glucose free media) and 6 days (labelled IR media) (A). The apoptosis pathway was negatively enriched and demonstrated reduced expression for insulin-resistant EHTs (B, C).

**Supplementary Figure 2.**
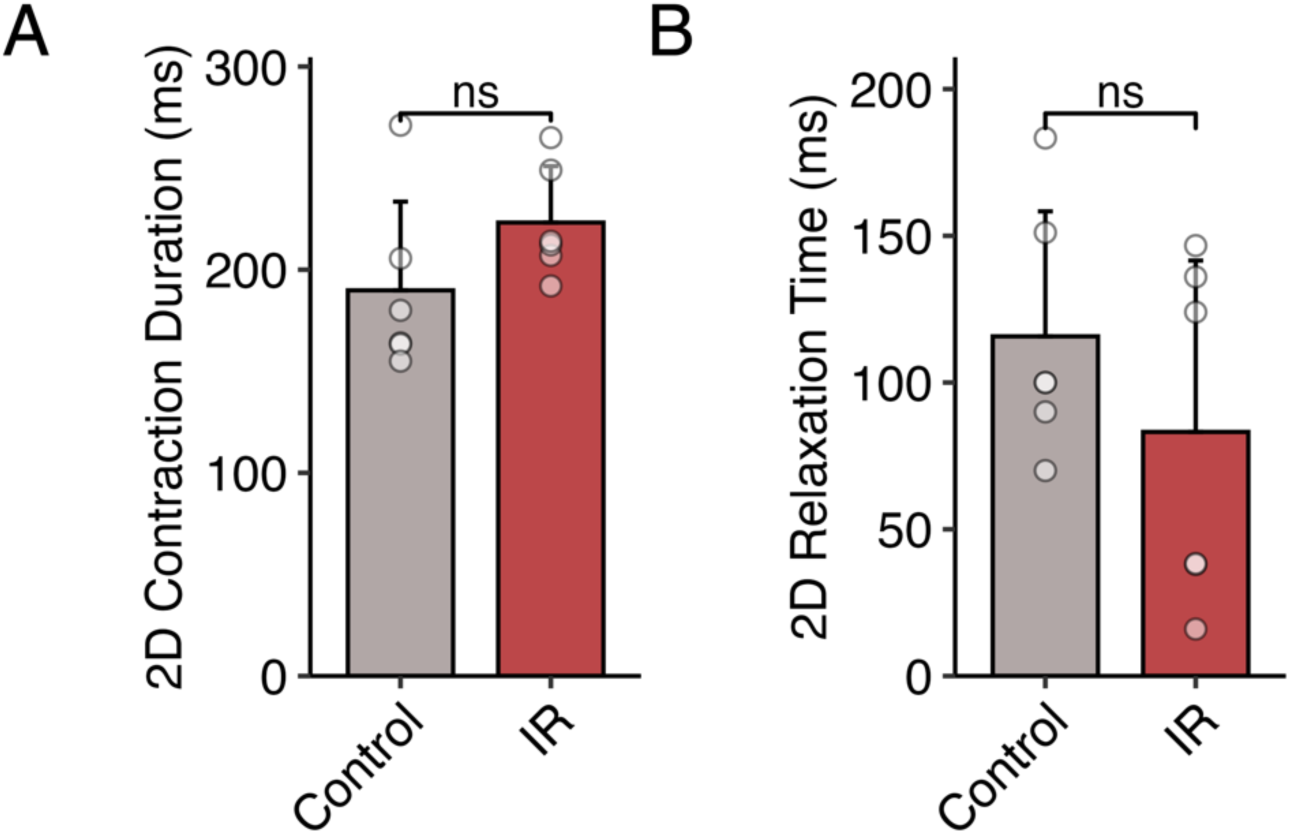
Culturing 2D hiPSC-CMs in IR media did not significantly change contraction duration and relaxation time. Musclemotion-based measurements of contraction and relaxation were not significantly altered between control and insulin resistant 2D hiPSC-CMs (A, B).

**Supplementary Table 1:**
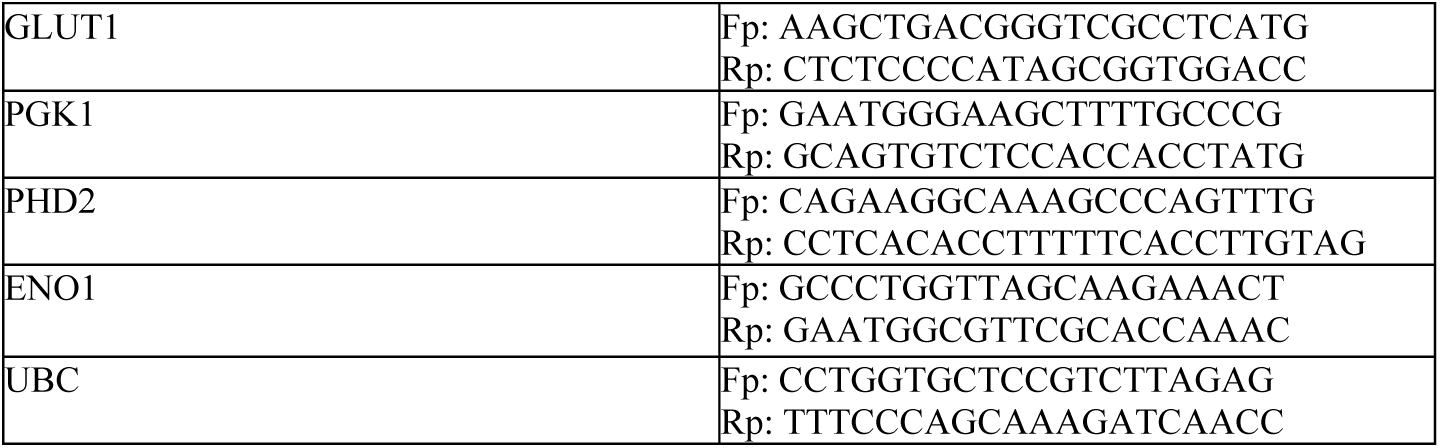
qPCR human primer sequences.

